# A *C. elegans* model of *C9orf72*-associated ALS/FTD uncovers a conserved role for eIF2D in RAN translation

**DOI:** 10.1101/2020.06.13.150029

**Authors:** Yoshifumi Sonobe, Jihad Aburas, Priota Islam, Tania F. Gendron, André E.X. Brown, Raymond P. Roos, Paschalis Kratsios

## Abstract

A hexanucleotide repeat expansion GGGGCC in the noncoding region of *C9orf72* is the most common cause of inherited amyotrophic lateral sclerosis (ALS) and frontotemporal dementia (FTD). Potentially toxic dipeptide repeats (DPRs) are synthesized from the GGGGCC sequence via repeat associated non-AUG (RAN) translation. We developed *C. elegans* models that express, either ubiquitously or exclusively in neurons, a transgene with 75 GGGGCC repeats flanked by intronic *C9orf72* sequence. The worms generate DPRs (poly-glycine-alanine [poly-GA], poly-glycine-proline [poly-GP]) and display neurodegeneration, locomotor and lifespan defects. Mutation of a non-canonical translation-initiating codon (CUG) upstream of the repeats blocked poly-GA production and ameliorated disease, suggesting poly-GA is pathogenic. Importantly, eukaryotic translation initiation factor 2D (*eif-2D/eIF2D*) was necessary for RAN translation. Genetic removal of *eif-2D* increased lifespan in both *C. elegans* models. *In vitro* findings in human cells demonstrated a conserved role for *eif-2D/eIF2D* in RAN translation that could be exploited for ALS and FTD therapy.

## INTRODUCTION

The GGGGCC (G4C2) hexanucleotide repeat expansion in the non-coding region of *C9orf72* is the most common monogenic cause of inherited amyotrophic lateral sclerosis (ALS) and frontotemporal dementia (FTD)^1, 2^, and also causes up to 10% of what appears to be sporadic ALS. The G4C2 repeat expands in patients to hundreds or thousands of copies that vary in number in different cells of the same individual. This genetic insult is thought to cause ALS/FTD pathogenesis via three non-mutually exclusive mechanisms: (1) loss of function due to decreased expression of C9ORF72 protein, (2) RNA toxicity from bidirectionally transcribed sense (GGGGCC) and anti-sense (CCCCGG) transcripts, and (3) proteotoxicity from dipeptide repeat (DPR) proteins produced from the expanded nucleotide repeats^3^.

This study focuses on the molecular mechanisms underlying DPR toxicity. Strong evidence suggests that DPRs are toxic in both cell culture and animal models of disease^4, 5^. Despite the presence of the expanded G4C2 repeat in the non-coding region of the RNA and the absence of an AUG initiating codon, DPRs are translated in all three reading frames from both sense and anti-sense transcripts through a process called repeat-associated non-AUG (RAN) translation^4^. Poly-glycine-alanine (poly-GA), poly-glycine-proline (poly-GP), and poly-glycine-arginine (poly-GR) are produced from sense transcripts, whereas poly-proline-arginine (poly-PR), poly-proline-alanine (poly-PA), and poly-GP are generated from antisense transcripts^6, 7, 8^. DPRs are present in neural cells of patients with *C9orf72*-associated ALS or FTD, indicating that RAN translation occurs *in vivo*^6, 7, 9, 10^. A number of potential causes for DPR toxicity have been proposed, including protein aggregation, impaired nucleocytoplasmic transport, and proteasome dysfunction^5^; however, the relative contribution of each DPR to disease pathogenesis remains unclear.

Recent *in vitro* and *in vivo* studies have identified non-canonical translation initiation factors as critical mediators of DPR production. For example, a recent study showed that the ribosomal protein eS25 (RPS25), which is required for efficient translation initiation at the internal ribosome entry site (IRES) of a number of viral and cellular RNAs^11, 12^, is also required for RAN translation of poly-GP from G4C2 repeats in yeast, fly, and induced pluripotent human stem cell models^13^. In addition, the eukaryotic translation initiation factor eIF3F, which is thought to control translation of hepatitis C viral (HCV) RNA through binding to the IRES of the viral genome^14^, regulates poly-GP production from G4C2 repeats in HEK293 cells^15^. Moreover, a fly study found that eIF4B and eIF4H are important for RAN translation of poly-GR from G4C2 transcripts^16^. Lastly, knockdown of eIF2A, an unconventional translation initiation factor used by cells under stress conditions^17, 18, 19^, had a partial role in poly-GA production in HEK293 cells and neural cells of the chick embryo^20^. Collectively, these and other studies^21, 22^ suggest that multiple factors are involved in RAN translation of DPRs from G4C2 transcripts; yet, their identity and molecular mechanisms of action remain elusive.

To identify novel factors involved in RAN translation of DPRs *in vivo*, we developed two *C. elegans* models for *C9orf72*-associated ALS/FTD. *C. elegans* provides a powerful system for the study of molecular and cellular mechanisms underlying neurodegenerative disorders due to: (1) its short lifespan (~3-4 weeks), (2) its compact nervous system (302 neurons in total), (3) the ability to study neuronal morphology and function with single-cell resolution, (4) the fact that most *C. elegans* genes have human orthologs, and (5) the ease of genomic engineering and transgenesis, which enables the rapid generation of worms that express human genes, permitting *in vivo* modeling of neurodegenerative disorders.

In this study, we initially developed *C. elegans* transgenic animals that carry, under the control of a ubiquitous promoter, 75 copies of the G4C2 repeat flanked by intronic *C9orf72* sequences. Compared to controls, these animals produce DPRs, namely poly-glycine-alanine (poly-GA) and poly-glycine-proline (poly-GP), and display evidence of neurodegeneration, as well as locomotor and lifespan defects. A second set of transgenic worms expressing the 75 G4C2 repeats exclusively in neurons displayed similar phenotypes, indicating that DPR production in this cell type is sufficient to cause disease phenotypes. These *C. elegans* models provide an opportunity to study *in vivo* the molecular mechanisms underlying DPR production. Through a candidate approach, we identified a novel role for the non-canonical translation initiation factor *eif-2D* (ortholog of human *eIF2D)* in RAN translation of G4C2 repeats. Genetic removal of *eif-2D* significantly decreased DPR production, and improved locomotor activity and lifespan in both *C. elegans* models. Supporting the phylogenetic conservation of our *C. ele*gans findings, knock-down of eIF2D in human cells led to decreased DPR production. Together, these observations suggest that eIF2D plays an important role in RAN translation of DPRs from G4C2 repeats, opening new directions for treatment of *C9orf72*-associated ALS and FTD.

## RESULTS

### Generation of a new *C. elegans* model for *C9orf72*-mediated ALS/FTD

The *C. elegans* genome contains a *C9orf72* ortholog (*alfa-1*), which is involved in lysosomal homeostasis, cellular metabolism and stress-induced neurodegeneration^23, 24, 25^. However, *alfa-1* does not contain G4C2 repeats. To study the molecular mechanisms underlying DPR production from G4C2 repeats, we generated worms that carry a transgene encoding 75 copies of the G4C2 sequence under the control of a ubiquitous (*snb-1*) promoter^26^ (**Fig. 1**A, see Materials and Methods). These 75 copies are flanked with intronic sequences normally found upstream (113 nucleotides) and downstream (99 nucleotides) of the G4C2 repeats in the human *C9orf72* gene (**Fig. 1**A). To monitor expression of poly-GA, the most amyloidogenic DPR^27, 28^, the transgene carries a nanoluciferase (nLuc) reporter in the poly-GA reading frame. Hereafter, we will refer to these transgenic animals as C9 ^*ubi*^ (**Fig. 1A**). In parallel, we generated two control *C. elegans* strains: (a) the UAG ^*ubi*^ worms carry an identical sequence to the C9 animals, but the upstream translation initiation codon CUG, which is required for translation of poly-GA *in vitro*^20, 29, 30, 31^, is mutated to UAG, and (b) the ΔC9 ^*ubi*^ worms lack the G4C2 repeats and intronic sequence of human *C9orf72* (**Fig. 1**A). Three independent transgenic lines for C9 ^*ubi*^, ΔC9 ^*ubi*^, and UAG ^*ubi*^ worms were generated and tested as described below.

**Figure 1.**
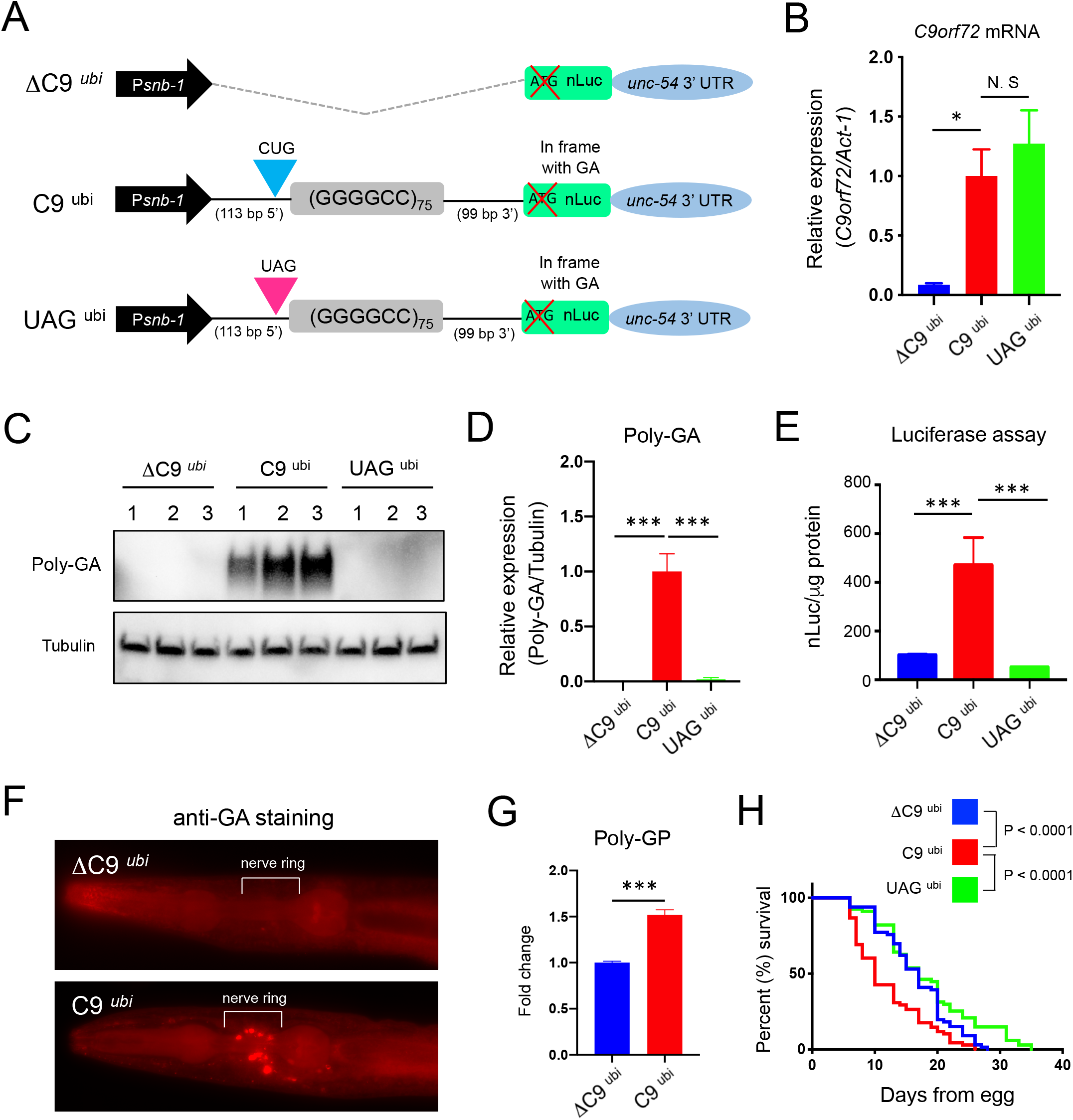
A *C. elegans* model for *C9orf72-mediated* ALS/FTD. (A) Schematic diagram showing ΔC9 ^*ubi*^, C9 ^*ubi*^, and UAG ^*ubi*^ constructs with the *snb-1* promoter. (B) The G4C2 repeat RNA (with *C9orf72* intronic RNA) and *act-1* mRNAs were assessed by RT-PCR (n = 3 different transgenic lines in each transgenic worm, mean ± s.e.m.). (C) Worm lysates were processed for Western blotting and immunostained with poly-GA and α-tubulin antibodies. (D) Quantification of poly-GA expression on Western blots (n = 3 different transgenic lines in each transgenic worm, mean ± s.e.m.) (E) Luciferae assay was performed in worm lysates (n = 3 different transgenic lines in each transgenic worm, mean ± s.e.m.). (F) Immunofluorescent staining for poly-GA (red) in ΔC9 ^*ubi*^ and C9 ^*ubi*^ adult worms. (G) Quantification of poly-GP in worms ΔC9 ^*ubi*^ (*kasIs8*) and C9 ^*ubi*^ (*kasIs7*) as determined by ELISA (n = 4, mean ± s.e.m.). (H) Lifespan of ΔC9 ^*ubi*^, C9 ^*ubi*^, or UAG ^*ubi*^ worms. Three different transgenic lines per genotype were used and the results are combined per genotype. ΔC9^*ubi*^: n = 66, C9^*ubi*^: n = 68, UAG^*ubi*^: n = 67). * P < 0.05, *** P < 0.001, N. S: not significant.

First, we performed quantitative PCR with reverse transcription (RT-qPCR) and found that the G4C2 repeats (*C9orf72* intronic mRNA) are only transcribed in C9 ^*ubi*^ and UAG ^*ubi*^ animals (**Fig. 1B**). Next, we tested whether DPRs (poly-GA, poly-GP, poly-GR) from the sense transcript (GGGGCC) are produced in the C9 ^*ubi*^ animals. We focused on these because the 75 G4C2 repeats are downstream of the *snb-1* promoter, suggesting that the sense transcript may be the one primarily transcribed and translated. Western blotting showed that poly-GA was robustly expressed in C9 ^*ubi*^ worms when compared to ΔC9 ^*ubi*^ and UAG ^*ubi*^ control animals (**Fig. 1**C, D). Importantly, mutating the non-canonical initiation codon CUG to UAG did not affect *C9orf72* mRNA levels, but led to a dramatic decrease in poly-GA expression (**Fig. 1**B-D). Similar results were obtained with a luciferase assay (**Fig. 1**E). Corroborating these observations, immunofluorescence staining revealed the presence of poly-GA aggregates in head neurons of C9 ^*ubi*^ animals (**Fig. 1**F). Although we did not detect poly-GP through Western blotting, a more sensitive ELISA assay detected a modest but statistically significant increase of poly-GP in the C9 ^ubi^ worms above the background level observed in ΔC9 ^*ubi*^ controls (**Fig. 1G**). These background levels likely reflect the abundance of GP motifs in the *C. elegans* proteome. Although we were not able to detect poly-GR in the C9 ^*ubi*^ worms (**Supplementary Figure 1A**), our overall results demonstrate that RAN translation occurs in *C. elegans*.

Importantly, we found that the median survival of C9 ^*ubi*^ worms is significantly shorter when compared to ΔC9 ^*ubi*^ and UAG ^*ubi*^ controls (P < 0.0001) (**Fig. 1**H), presumably due to poly-GA- and/or poly-GP-associated toxicity. Together, these findings suggest that at least two distinct DPRs (poly-GA and poly-GP) are generated from the 75 G4C2 repeats in the C9 ^*ubi*^ animals and likely contribute to the reduced survival of these animals, thereby establishing a new *C. elegans* model for *C9orf72-mediated* ALS/FTD.

### C9 ^*ubi*^ animals display progressive motor neuron degeneration and neuromuscular synapse defects

At the level of animal behavior, we observed striking locomotion defects in C9 ^*ubi*^ worms, which are presented later in Results. Such defects could be attributed, at least partially, to degeneration of ventral nerve cord motor neurons, which are essential for *C. elegans* locomotion. To test this possibility, we fluorescently labeled the cholinergic motor neurons present in the nerve cord of C9 ^ubi^ and ΔC9 ^ubi^ animals with the *unc-17/VAChT::gfp* transgenic reporter, which allows for single-cell resolution in our analysis (**Fig. 2**A-B). Next, we quantified the number of cholinergic motor neurons (*unc-17/VAChT::gfp* positive) in C9 ^*ubi*^ and ΔC9 ^*ubi*^ animals at juvenile (larval stage 4, L4) and adult (Day 2 and 5) stages. Although no differences were observed at the juvenile stage, we found a significant reduction in the number of motor neurons that populate the nerve cord of C9 ^*ubi*^ animals (**Fig. 2**C), arguing for progressive motor neuron degeneration. Of note, we cannot exclude the possibility that, in addition to cholinergic motor neurons, other neuron types also degenerate in C9 ^*ubi*^ animals.

**Figure 2.**
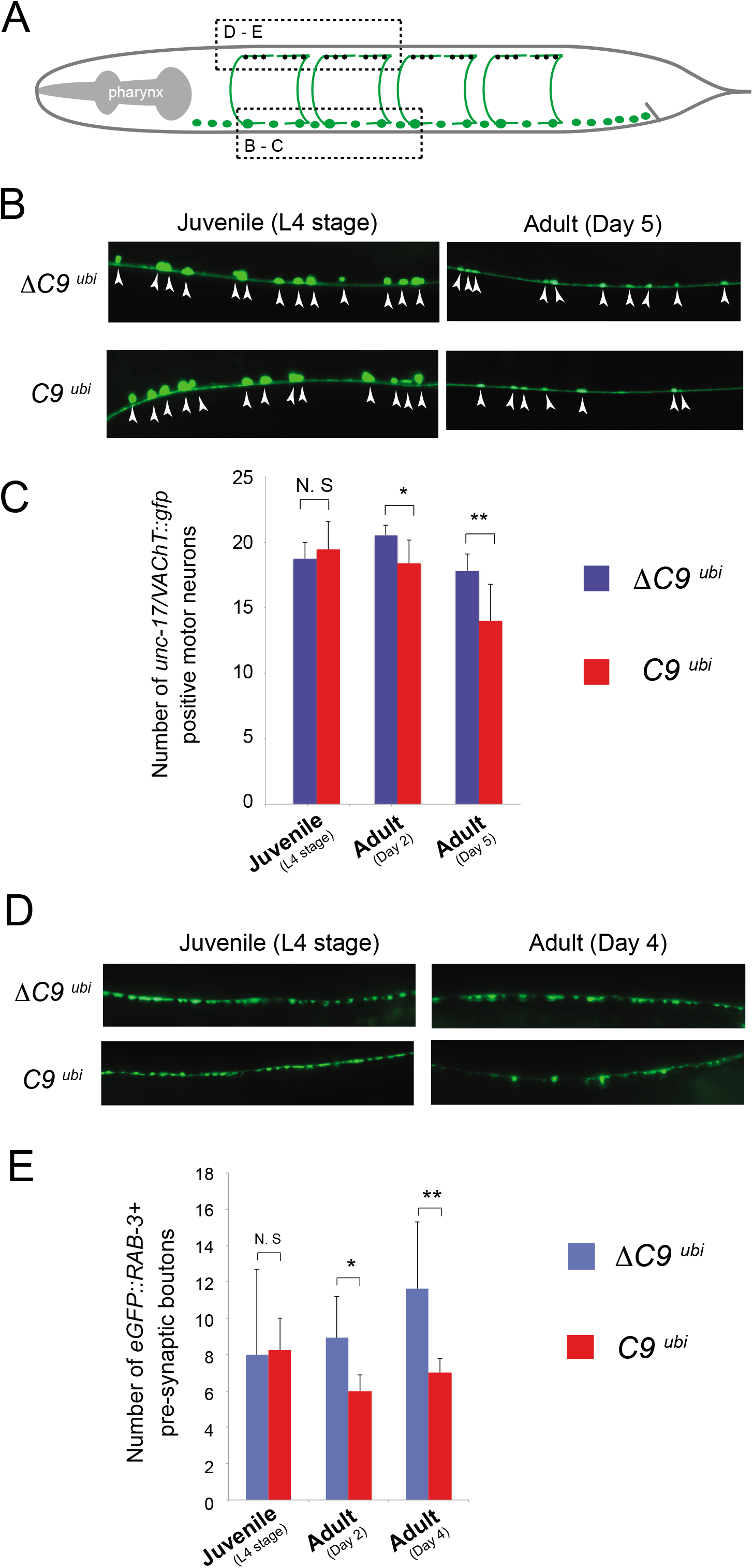
Evidence for motor neuron degeneration and synaptic defects in C9 ^*ubi*^ animals. (A) Schematic showing cholinergic motor neurons (green) in the *C. elegans* ventral nerve cord. Dashed boxes indicate the regions analyzed in panels B-C and C-D. (B) Motor neuron cell bodies are visualized with the *vsIs48* [*unc-17/VAChT::gfp*] marker in ΔC9 ^*ubi*^ and C9 ^*ubi*^ animals at the indicated stages. Arrowheads indicate motor neuron cell bodies. (C) Quantification of panel B. N >10 per genotype. *: p <0.05; **: p < 0.01; N.S: not statistically significant. (D) The presynaptic boutons of cholinergic motor neurons on dorsal body wall muscle are visualized with *otIs437 [Punc-3::eGFP::rab-3cDNA]* in ΔC9 ^*ubi*^ and C9 ^*ubi*^ animals at the indicated stages. (E) Quantification of panel D. N >10 per genotype. *: p <0.05; **: p < 0.01; N.S: not statistically significant.

We next sought to determine whether the observed motor neuron degeneration is accompanied by neuromuscular synapse defects, an early hallmark in ALS disease progression^32^. To this end, we crossed the C9 ^*ubi*^ worms to animals carrying a fluorescent reporter (*Punc-3::rab-3::eGFP*) that specifically labels the presynaptic boutons of cholinergic motor neurons (**Fig. 2**D). We found that the number of presynaptic boutons is significantly decreased in C9 ^*ubi*^ animals during adult (Day 2 and 4) stages, but no differences were observed between C9 ^*ubi*^ and ΔC9 ^*ubi*^ animals at a juvenile stage (L4) (**Fig. 2**E). In summary, these findings provide evidence for motor neuron degeneration and neuromuscular synapse defects in C9 ^*ubi*^ animals.

### DPR production selectively in neurons is sufficient to cause disease phenotypes *in vivo*

The DPRs (poly-GA, poly-GP) in *C9 ^ubi^* animals are produced ubiquitously because the *snb-1* promoter is active in all *C. elegans* cell types (**Fig. 1**A)^26^. To test whether DPR production from the G4C2 repeats exclusively in neurons is sufficient to cause disease-associated phenotypes, we generated a second set of transgenic animals by replacing the *snb-1* promoter with a fragment of the *unc-11* promoter (*unc-11c*) known to be exclusively active in all *C. elegans* neurons^26^. We refer to these animals as C9 ^*neuro*^, ΔC9 ^*neuro*^, and UAG ^*neuro*^ (**Fig. 3**A). First, we confirmed by RT-qPCR that the G4C2 repeat RNA (with *C9orf72* intronic mRNA) and the *nLuc* mRNA are transcribed in C9 ^*neuro*^ and UAG ^*neuro*^ animals (**Fig. 3**B-C). Similar to C9 ^*ubi*^ animals, we found through Western blotting that poly-GA is robustly produced in C9 ^*neuro*^ animals, but not in control animals lacking the G4C2 repeats (ΔC9 ^*neuro*^) or having the non-canonical translation initiation codon CUG mutated to UAG (UAG ^*neuro*)^ (**Fig. 3**D-E). However, we were not able to detect poly-GR by ELISA in the C9 ^*neuro*^ animals (**Supplementary Figure 1B**). Importantly, we found a dramatic decrease in the median survival of C9 ^*neuro*^ animals when compared to ΔC9 ^*neuro*^, and UAG ^*neuro*^ controls (p value < 0.0001) (**Fig. 3**F). The effect on median survival in C9 ^*neuro*^ animals appears stronger when compared to C9 ^*ubi*^ animals (**Fig. 1**H, 3F); this is likely due to differential strengths of the *snb-1* and *unc-11c* promoters, resulting in different amounts of DPRs. Our findings suggest that DPR (poly-GA) production from the G4C2 repeats exclusively in neurons is sufficient to trigger disease-associated phenotypes, establishing C9 ^*neuro*^ animals as a model for *C9orf72*-mediated ALS/FTD.

**Figure 3.**
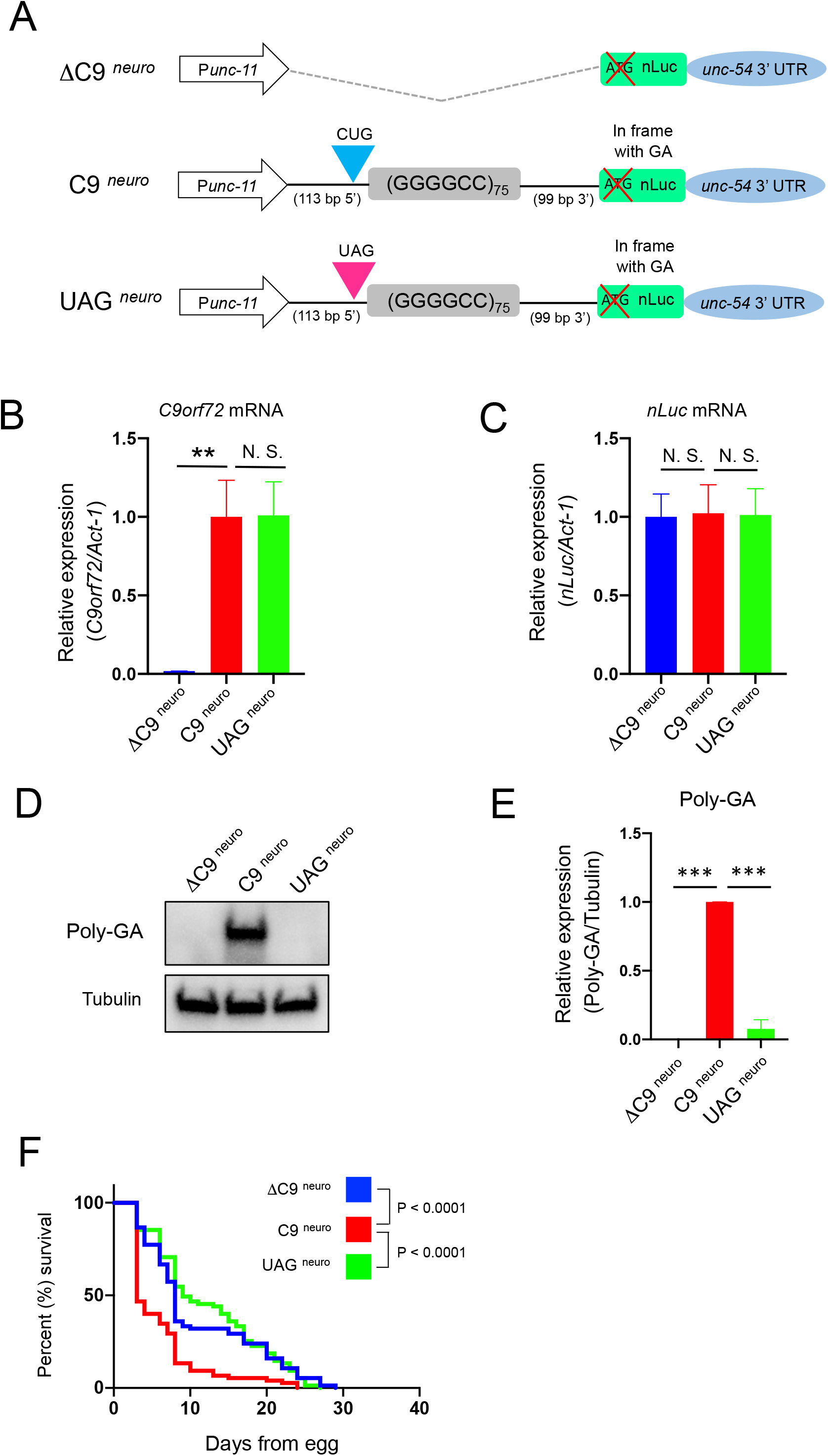
DPR expression exclusively in neurons reduces *C. elegans* lifespan. (A) Schematic diagram showing ΔC9^*neuro*^(kasEx159), C9^*neuro*^ (kasEx157), and UAG^*neuro*^ (kasEx160) worms under the *unc11c* promoter. (B) The G4C2 repeat RNA (with *C9orf72* intronic RNA) and act-1 mRNAs were assessed by RT-PCR. (C) The nLuc and act-1 mRNAs were assessed by RT-PCR (The experiments were repeated 4 times using 1 transgenic line per genotype, mean ± s.e.m.). (D) Worm lysates were processed for Western blotting and immunostained with poly-GA and tubulin antibodies. (E) Quantification of poly-GA expression on Western blots. The experiments were repeated 3 times using 1 transgenic line per genotype, mean ± s.e.m.) (F) Lifespan of ΔC9*^neuro^* (kasEx159), C9^*neuro*^ (kasEx157), or UAG^*neuro*^ (kasEx160) worms. The experiments were repeated 3 times using 1 transgenic line per genotype, mean ± s.e.m. ΔC9^*neuro*^ (kasEx159): n = 75, C9^*neuro*^ (kasEx157): n = 75, UAG^*neuro*^ (kasEx160): n = 75). ** P < 0.01, **** P < 0.0001, n.s. not significant.

### Genetic removal of the eukaryotic translation initiation factor 2D (*eif-2D/eIF2D*) reduces poly-GA production and ameliorates lifespan defects in C9 ^*ubi*^ and C9 ^*neuro*^ animals

The C9 ^*ubi*^ and C9 ^*neuro*^ worm models for *C9orf72*-mediated ALS/FTD offer a unique opportunity to uncover the molecular mechanisms responsible for RAN translated DPRs from G4C2 repeats. Our results strongly suggest that RAN (non-AUG dependent) translation of poly-GA in both *in vivo* models (C9 ^*ubi*^ and C9 ^*neuro*^ worms) requires the non-canonical initiation codon CUG. We therefore surveyed the literature for factors known to initiate translation via this codon. Two proteins, the eukaryotic translation initiation factors *eIF2A* and *eIF2D*, are known to deliver tRNAs to the CUG codon and initiate translation *in vitro*^33, 34^. A single ortholog for each factor is present in the *C. elegans* genome, that is, *eif-2A* for human *eIF2A* and *eif-2D* for human *eIF2D*. We therefore questioned whether *eif-2A* and/or *eif-2D* are necessary for RAN translation of DPRs from the G4C2 repeats. To test this, we obtained mutant animals that carry strong loss-of-function (putative null) alleles for *eif-2A* (*gk358198*) or *eif-2D* (*gk904876*). Since these animals are viable and fertile, we crossed them to the C9 ^ubi^ worms and generated *eif-2A* (*gk358198*); C9 ^*ubi*^ and *eif-2D* (*gk904876*); C9 ^*ubi*^ animals. Next, we found via RT-qPCR that the G4C2 repeat RNA with flanking *C9orf72* intronic sequence, as well as the *nLuc* mRNA are normally transcribed in both *eif-2A* (*gk358198*); C9 ^*ubi*^ and *eif-2D* (*gk904876*); C9 ^*ubi*^ animals (**Fig. 4**A-B), indicating that loss of *eif-2A* or *eif-2D* does not affect transcription or stability of the repeat RNA. However, we found a significant decrease in DPR (poly-GA) production in *eif-2D* (*gk904876*); C9 ^*ubi*^ animals but not in *eif-2A* (*gk358198*); C9 ^*ubi*^ animals (**Fig. 4**C-D), suggesting that *eif-2D* is required for RAN translation. *eif-2D* does not appear to play a role in canonical translation because loss of *eif-2D* gene activity in worms did not affect expression of an AUG-initiated green fluorescent protein (GFP) transgene (**Supplementary Figure 2**).

**Figure 4.**
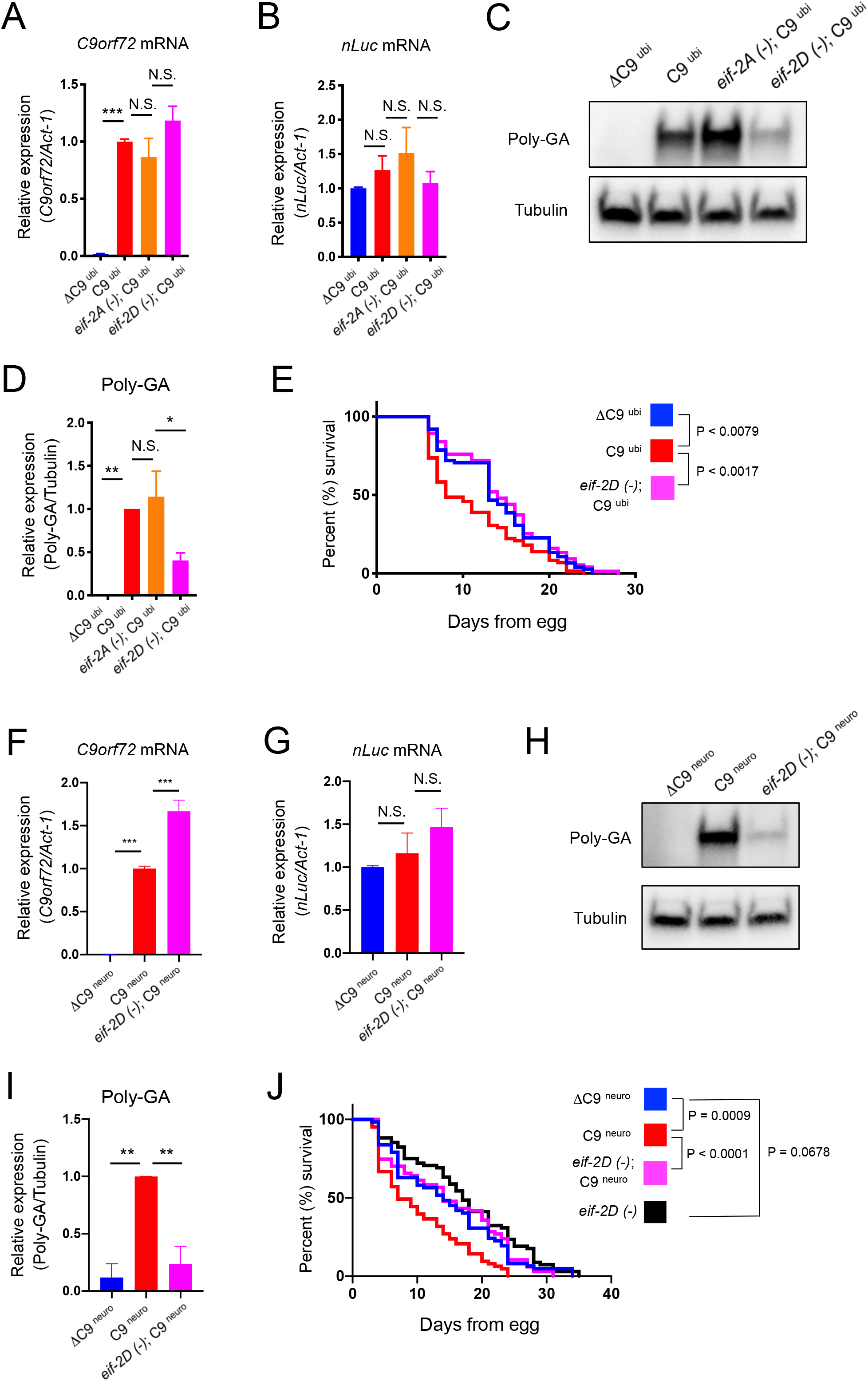
Genetic removal of *eif-2D/eIF2D* in C9 ^*ubi*^ and C9 ^*neuro*^ animals suppresses poly-GA expression and ameliorates lifespan defects. (A and F) The G4C2 repeat RNA (with *C9orf72* intronic RNA) and Act-1 mRNAs were assessed by RT-PCR. (B and G) The nLuc and Act-1 mRNAs were assessed by RT-PCR. The experiments were repeated 4 times in C and 3 times in G, respectively, using 1 transgenic line per genotype (mean ± s.e.m). (C and H) Worm lysates were processed for Western blotting and immunostained with poly-GA and tubulin antibodies. (D and I) Quantification of poly-GA expression on Western blots. The experiments were repeated 4 times in B and 3 times in F, respectively, using 1 transgenic line per genotype (mean ± s.e.m.) (E and J) Lifespan of ΔC9^*ubi*^ (kasEx154), C9^*ubi*^, and *eif-2D* (*gk904876*); C9 ^*ubi*^ animals. One transgenic line was used per genotype. ΔC9^*ubi*^ (kasEx154): n = 75, C9^*ubi*^: n = 72, *eif-2D* (*gk904876*); C9 ^*ubi*^: n = 75) and ΔC9^*neuro*^ (kasEx159), C9^*neuro*^ (kasEx157), *eif-2D* (*gk904876*); C9^*neuro*^ (kasEx157), and *eif-2D* (*gk904876*) worms. One transgenic line was used per genotype. ΔC9^*neuro*^: n = 62, C9^*neuro*^: n = 63, *eif-2D* (*gk904876*); C9^*neuro*^: n = 67, *eif-2D* (*gk904876*): n = 68). * P < 0.05, ** P < 0.01, **** P < 0.0001.

Since *eif-2D* is necessary for translation of poly-GA, which is thought to be toxic, loss of *eif-2D* gene activity may improve survival in C9 ^*ubi*^ animals. Indeed, we found that the *eif-2D* (*gk904876*); C9 ^*ubi*^ animals display a normal lifespan, similar to ΔC9 ^*ubi*^ animals (**Fig. 4**E). Eukaryotic translation initiation factors are expressed broadly in many tissues, and this appears to be the case for *eif-2D* in *C. elegans (https://bgee.org/?page=gene&gene_id=WBGene00016113)* and its human ortholog *(https://www.proteinatlas.org/ENSG00000143486-EIF2D/tissue)*. To test whether neuronal *eif-2D* gene activity is critical for poly-GA production and animal survival, we crossed the *eif-2D (gk904876)* mutants to the C9 ^*neuro*^ animals that selectively produce poly-GA in neurons (**Fig. 3**D). Although transcription of the G4C2 repeats (*C9orf72* intronic mRNA) increased in *eif-2D* (*gk904876*); C9 ^*neuro*^ animals for unclear reasons (**Fig. 3**F-G), we found a dramatic decrease in poly-GA production in these worms compared to C9 ^*neuro*^ worms (**Fig. 3**H-I). Of note, genetic removal of *eif-2D* in *C9 ^neuro^* animals resulted in a more prominent decrease in poly-GA levels than in *C9 ^ubi^* animals (compare **Fig. 4**C-D with **Fig. 4**H-I), suggesting that eIF2D may have a greater impact on RAN translation in neurons than in other cell types. Importantly, loss of *eif-2D* restored survival of C9 ^*neuro*^ animals (P value < 0.0001 between *eif-2D* (*gk904876*); C9 ^*neuro*^ and C9 ^*neuro*^) to levels similar to ΔC9 ^*neuro*^ and *eif-2D* (*gk904876*) control animals (**Fig. 3**J). In summary, we found that *eif-2D/eIF2D* is critical for poly-GA production, and its genetic removal mitigates the shortened lifespan otherwise observed in both C9 ^*ubi*^ and C9 ^*neuro*^ animals.

### Genetic removal of *eif-2D/eIF2D* ameliorates the locomotor defects of *C9 ^ubi^* animals

Because loss of *eif-2D* reduces poly-GA production and improves animal survival (**Fig. 4**), we next asked whether the locomotor defects of C9 ^*ubi*^ animals can be ameliorated upon genetic removal of *eif-2D*. To this end, we performed a high-resolution behavioral analysis of freely moving adult (day 2) *C. elegans* using automated worm tracking technology^35, 36^. This analysis can quantitate several features related to locomotion (*e.g*., velocity, crawling amplitude, body curvature). We found multiple locomotor defects in C9 ^*ubi*^ animals, such as reduced velocity and impaired body curvature (**Fig. 5, Supplementary Figure 3**). These defects were not present in ΔC9 ^ubi^ and UAG ^ubi^ controls indicating that a hallmark of ALS, namely motor deficits, is recapitulated in the C9 ^*ubi*^ animal model. Importantly, we performed the same tracking analysis on *eif-2D* (*gk904876*) mutants and *eif-2D* (*gk904876*); C9 ^*ubi*^ animals. This analysis revealed that loss of *eif-2D* significantly improved the locomotor defects of C9 ^*ubi*^ animals (**Fig. 5, Supplementary Figure 3**). Of note, *eif-2D* (*gk904876*) single mutants did not display any prominent locomotor defects (**Fig. 5**). In summary, this behavioral analysis complements our molecular analysis (poly-GA detection) and survival assays (**Fig. 4**), providing compelling evidence that loss of *eif-2D* ameliorates the pathogenic phenotype associated with RAN translation of *C9orf72* G4C2 nucleotide repeats.

**Figure 5.**
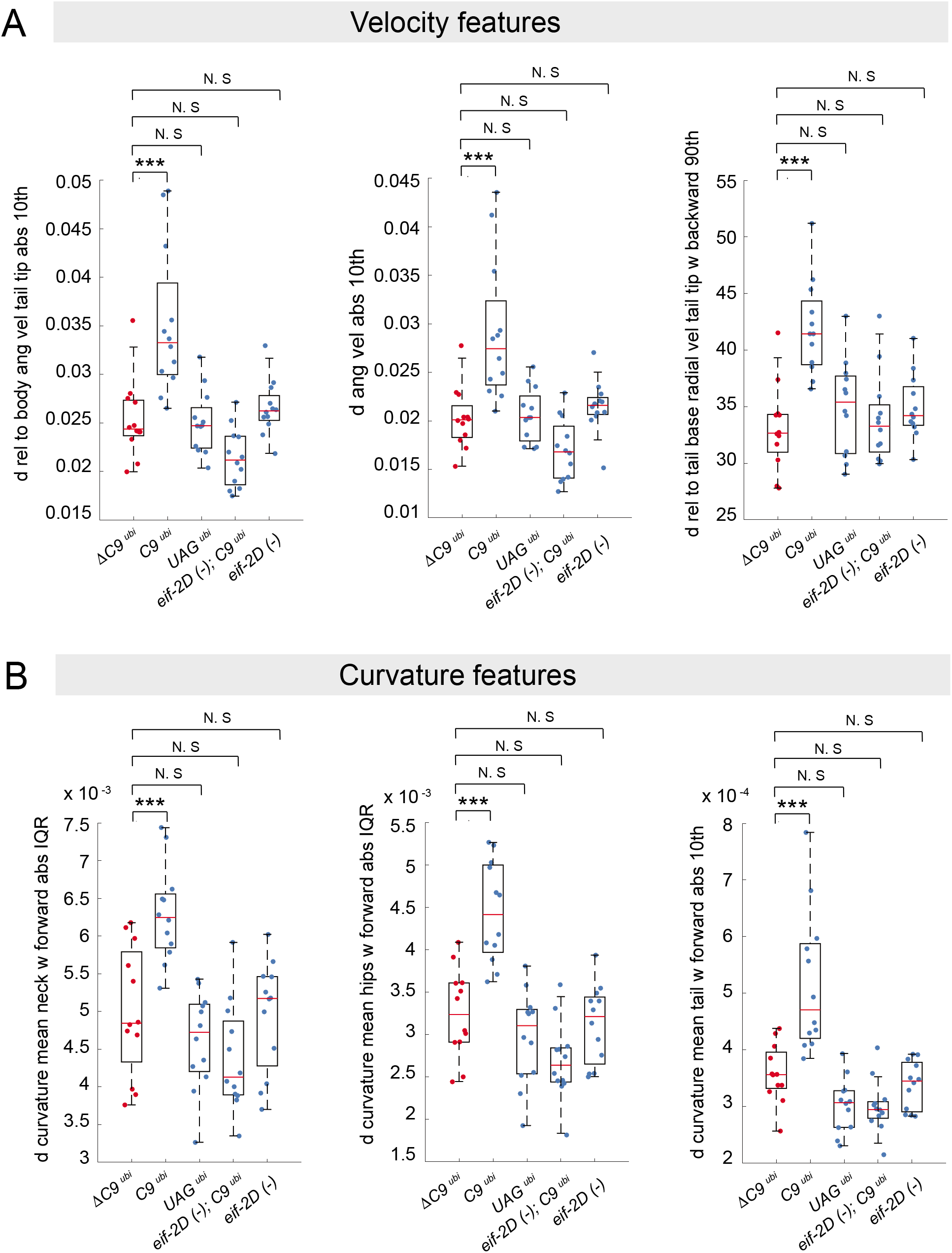
Genetic removal of *eif-2D/eIF2D* ameliorates the locomotor defects of *C9 ^ubi^* animals. Examples of locomotion features significantly affected in *C9 ^ubi^* animals. Genetic removal of *eif-2D* ameliorated the locomotion defects of *C9 ^ubi^* animals [*eif-2D (-); C9 ^ubi^*]. Tracking analysis was performed at Day 2 adult animals (N = 12). The p-value threshold for this analysis (FDR 5%) is 0.0014 (see Methods). Panel A shows three locomotion features related to animal velocity. *d rel to body ang vel tail tip abs 10th*: 10th percentile of the derivative of the absolute value of the angular velocity of the tip of the tail relative to the centroid of the mid-body points. *d ang vel abs 10^th^*: 10th percentile of the derivative of the absolute value of the angular velocity of the worm. *d rel to tail base radial vel tail tip w backward 90^th^*: 90th percentile of the derivative of the radial velocity of the tip of the tail relative to the centroid of the tail base points, while the worm is moving backwards. Panel B shows three locomotion features related to body curvature while the worm is moving forward. *d curvature mean neck w forward abs IQR*: interquartile range of the derivative of the absolute value of the mean curvature of the neck. *d curvature mean hips w forward abs IQR*: interquartile range of the derivative of the absolute value of the mean curvature of the hips. *d curvature mean tail w forward abs 10^th^*: 10th percentile of the derivative of the absolute value of the mean curvature of the tail.

### Knock-down of *eIF2D* suppresses DPR production in human cells

We next sought to determine whether the function of *eif-2D/eIF2D* in RAN translation and DPR production is conserved from worms to human cells. To this end, we transfected HEK293 cells with a bicistronic construct, in which the firefly luciferase (fLuc) gene is located in the first cistron followed by a second cistron containing the 75 G4C2 repeats along with intronic sequences that normally flank the repeats in the human *C9orf72* locus (**Fig. 6**A). To monitor expression of different DPRs, we generated three different versions of this construct by inserting a nanoluciferase (nLuc) reporter gene in the poly-GA, poly-GP, or poly-GR reading frame (**Fig. 6**A) (see Materials and Methods). Importantly, previous studies showed that DPR translation can occur when the G4C2 repeats are located in the second cistron of a bicistronic construct^20, 37^. Each bicistronic construct was co-transfected with plasmids carrying either control short hairpin RNA (shRNA) or shRNA against *eIF2D*. First, we assessed the eIF2D protein levels via Western blotting and observed a close to 50% reduction in HEK293 cells that received the *eIF2D* shRNA compared to cells transfected with control shRNA (**Fig. 6**B-C). Next, we performed Western blotting and luciferase assays and found that, similar to our *C. elegans* model, poly-GA and poly-GP peptides are produced in HEK293 cells upon transfection (**Fig. 6**B, D-E). Moreover, the increased sensitivity of the luciferase assay enabled us to detect poly-GR in these cells, albeit at lower levels compared to poly-GA (**Fig. 6**E). Importantly, knock-down of eIF2D by shRNA dramatically reduced (by at least 75%) the expression of all three DPRs (poly-GA, poly-GP, poly-GR), suggesting eIF2D is required for RAN translation of GA, GP, and GR (**Fig. 6**B, D-E). In contrast, the protein levels of fLuc, which is expressed by the first cistron via canonical AUG-dependent translation, were similar in cells transfected with either control or eIF2D shRNAs (**Fig. 6**B, F), suggesting a specific role for eIF2D in unconventional translation in human cells. In summary, these results suggest that eIF2D is necessary for DPR production (poly-GA, poly-GP, poly-GR) in human cultured cells, uncovering a conserved role for eIF2D in RAN translation of DPRs from G4C2 repeat-containing RNA.

**Figure 6.**
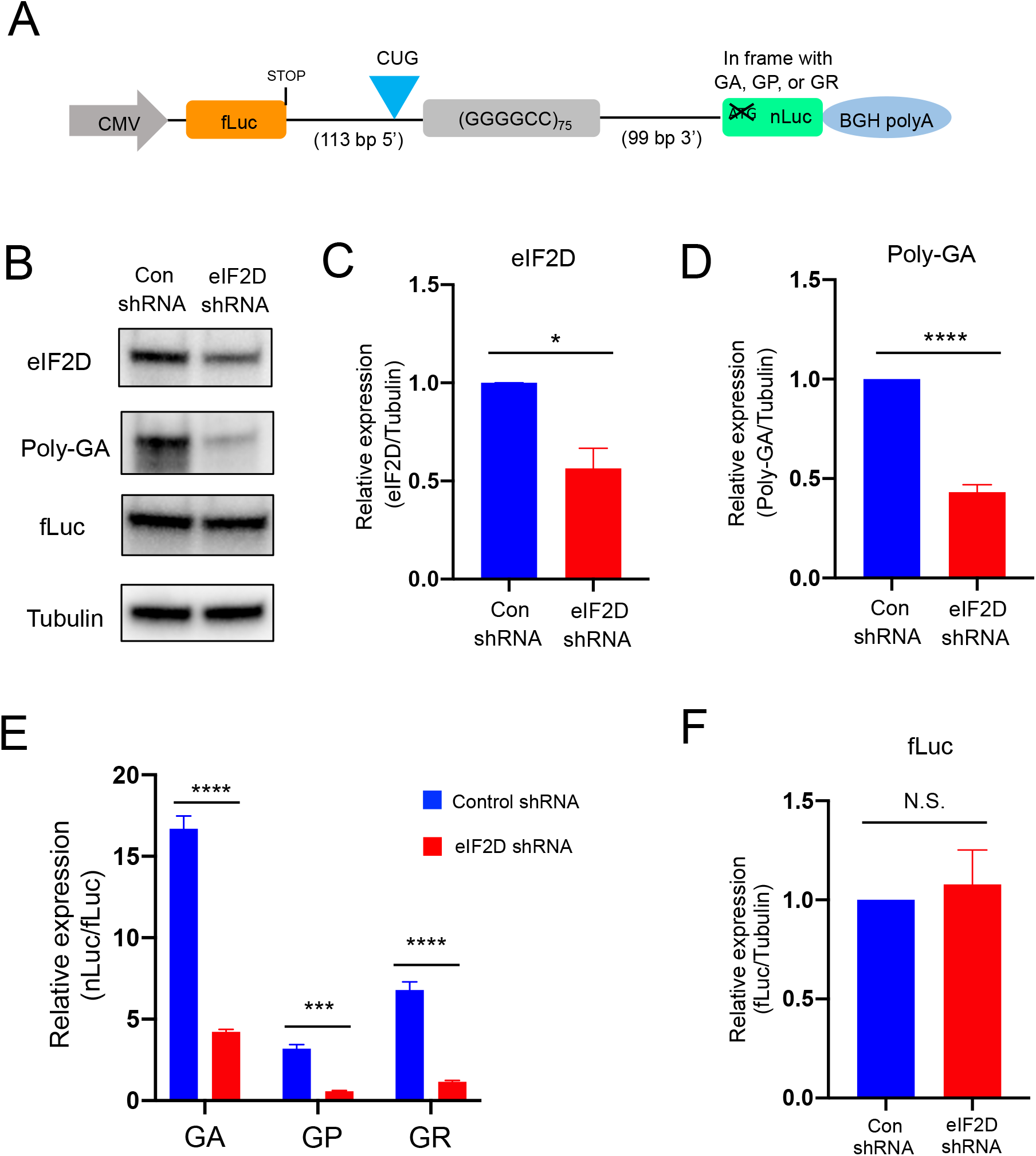
Knockdown of eIF2D suppresses DPR expression in HEK293 cells. Bicistronic constructs were cotransfected along with either control or anti-eIF2D shRNA and cultured for 48h. (A) Schematic diagram showing bicistronic constructs. (B) The cell lysates were processed for Western blotting and immunostained with poly-GA, fLuc, eIF2D, and tubulin antibodies. (C, D, and F) Quantification of (C) eIF2D, (D) poly-GA, and (F) fLuc on Western blots (The experiments were repeated 3 times. mean ± s.e.m.) (E) The levels of luciferase activity was assessed by dual luciferase assays (The experiments were repeated 3 times. mean ± s.e.m.) * P < 0.05, *** P < 0.001, **** P < 0.0001, n.s. not significant.

## DISCUSSION

The discovery of RAN translation of *C9orf72* nucleotide repeats combined with the reported toxicity of DPRs in model systems have focused the ALS/FTD field on this unconventional form of translation, as well as on DPRs as potential therapeutic targets. To study the molecular mechanisms underlying RAN translation, we initially developed a *C. elegans* model that carries a transgene encoding 75 copies of the G4C2 repeat under the control of a ubiquitous promoter. These C9 ^ubi^ animals generate DPRs, namely poly-GA and poly-GP, show signs of neurodegeneration, and display locomotor and lifespan defects. Such phenotypes are likely due to DPR toxicity because mutation of a non-canonical CUG initiation codon decreased poly-GA production and ameliorated locomotor and lifespan defects in C9 ^ubi^ animals without affecting *C9orf72* mRNA levels. Moreover, a second *C. elegans* model (referred to as C9 ^neuro^) that generates poly-GA exclusively in neurons displayed a similar phenotype. Importantly, we found that knock-out of the eukaryotic translation initiation factor *eif-2D/eIF2D* suppressed poly-GA expression and ameliorated lifespan defects in both C9 ^ubi^ and C9 ^neuro^ models. Lastly, our *in vitro* findings in human cells combined with our *C. elegans* data argue for a conserved role of *eif-2D/eIF2D* in RAN translation that could open up new therapeutic directions for ALS and FTD.

### The C9 ^ubi^ and C9 ^neuro^ worm models offer insights into DPR toxicity

An unresolved issue in the field of ALS and FTD research is the relative contribution of each of the five DPRs (poly-GA, poly-GP, poly-GR, poly-PR, poly-PA) to disease pathogenesis. In this paper, two lines of evidence indicate that poly-GA, the most readily visible DPR in the brain and spinal cord of ALS/FTD patients with *C9orf72* expanded repeats^38, 39^, significantly contributes to disease phenotypes in *C. elegans:* (1) poly-GA aggregates and appears to be the most abundantly expressed DPR in our worm models, and (2) mutation of the non-canonical CUG initiation codon decreased poly-GA production and ameliorated locomotor and lifespan defects in C9 ^ubi^ and C9 ^neuro^ animals. Importantly, these *in vivo* results significantly extend previous reports demonstrating that this same CUG is required *in vitro* for poly-GA translation^20, 29, 30, 31^. In support of our *C. elegans* findings, recent studies in mice reported neurodegeneration, motor deficits, and cognitive defects following ubiquitous expression of poly-GA^40^, as well as behavioral abnormalities following spinal cord expression of poly-GA^41^. A prevailing hypothesis is that poly-GA causes cellular toxicity through proteosomal impairment and protein sequestration^5^. A recent publication reports that in an *in vitro* system poly-GA spreads cell-to-cell and inhibits proteasomal function both cell-autonomously and non-cell-autonomously, leading to ubiquitination of the nuclear localization signal of TDP-43 and its subsequent cytoplasmic mislocalization^42^. Importantly, treatment with an antibody against poly-GA ameliorated TDP-43 mislocalization^42^, as well as toxicity in mouse models for *C9orf72*-associated ALS/FTD^43^. However, a zebrafish model that produces poly-GA did not exhibit motor axon toxicity^44^, presumably due to low poly-GA expression.

Although our C9 ^ubi^ worm model produces a small amount of poly-GP, this DPR is thought to be relatively nontoxic^5^. In contrast, the toxicity of arginine-containing DPRs, especially poly-GR, has been demonstrated in mouse neurons^45^. In *C. elegans*, codon-optimized constructs driving expression of poly-GR or poly-PR caused toxicity in muscles and neurons^46^. Furthermore, initiation of poly-GR translation has been reported to occur at the same upstream CUG codon used for poly-GA production, but with subsequent frameshifting into the GR reading frame^30^. However, two recent papers showed that translation initiation of poly-GR did not occur from the upstream CUG codon^31, 47^. Although we could not detect poly-GR by Western blotting and ELISA (**Supplementary Figure 1**), it is possible that small amounts of poly-GR are produced in our *C. elegans* models and contribute to the observed disease phenotypes. Supporting this notion, very low levels of poly-GR in neurons have been reported to impair synaptic function and induce behavioral deficits in both *Drosophila* and mice^45^.

In addition to the possibility of DPR toxicity in our *C. elegans* models, there remains the question as to whether the repeat RNA itself rather than, or in addition to, its protein product could be toxic^48^. A role for RNA toxicity does not appear to be the case in our nematode experiments since mutation of the non-canonical initiation codon CUG to UAG led to a significant decrease in poly-GA expression and rescue of worm survival to wild type levels, but had no effect on *C9orf72* mRNA levels. However, it is possible that the CUG > UAG mutation, albeit minimal in nature, can affect RNA structure.

One question that arises is whether our transgenic C9 ^ubi^ and C9 ^neuro^ animals constitute an appropriate model for the *C9orf72* patient. It seems clear that the RNA template (repeat RNA plus flanking intronic sequence) produced in these animals differs from the RNA template actually expressed in the central nervous system of patients with a *C9orf72* expanded repeat because it is likely that there are varying RNA templates of different lengths and sequences in these patients. If this is indeed the case, it suggests that a number of potential toxicities besides one related to poly-GA production may be present in the human disease state, including toxicity related to repeat RNA and/or synthesis of other DPRs. Toxicity may also occur as a consequence of a decrease in the authentic *C9orf72* gene product. Supporting this possibility, a recent study reported that a decrease in C9ORF72 protein leads to a decrease in autophagic clearance of toxic accumulations of the DPRs^31^.

### A novel role for *eif-2D/eIF2D* in RAN translation of *C9orf72*-associated nucleotide repeats

eIF2D is known to initiate translation *in vitro* at both AUG and non-AUG initiation codons^33, 49^. The present study demonstrates a key role of eIF2D in RAN translation of poly-GA since eIF2D knock-down in HEK293 cells, as well as *eif-2D* knock-out in *C. elegans* led to a prominent decline in poly-GA expression. It is noteworthy that genetic removal of *eif-2D* in the worm had no effect on: (a) canonical translation of an AUG-initiated GFP reporter, (b) animal locomotion, and (c) lifespan. The absence of detrimental phenotypes in *C. elegans* animals that globally lack *eif-2D*/eIF2D make this factor an attractive therapeutic target. Moreover, we found that removal of *eif-2D* led to a greater decrease in DPR (poly-GA) production in C9 ^neuro^ animals versus C9 ^ubi^ animals (**Figure 4D, I**), suggesting that eIF2D may be regulating RAN translation in varying ways in different cell types.

Our previous study demonstrated that knock-down of eIF2A in HEK293 cells and neural cells in the chick embryo led to a modest (30%) decrease in poly-GA expression from G4C2 expanded repeats^20^. In the present study, however, *eif-2A/eIF2A* deficiency had no effect on poly-GA expression in *C. elegans*, while eIF2D knock-down in HEK293 cells and *eif-2D* knock-out in *C. elegans* had a prominent effect on poly-GA expression, suggesting eIF2D plays a more important role in RAN translation than eIF2A. In contrast to the slight decrease of a single DPR (poly-GA) we previously found with eIF2A knock-down^20^, knock-down of eIF2D in HEK293 cells dramatically reduced (by at least 75%) the expression of multiple DPRs (poly-GA, poly-GP and poly-GR), indicating a critical role for eIF2D in RAN translation in all three reading frames. It is possible, however, that eIF2A does play a significant role in RAN translation, but only upon cellular stress since it can translate a subset of mRNAs when eIF2α is phosphorylated^17, 18, 50^.

In addition to eIF2D, RPS25 is the only other factor known to be required for the expression of these three DPRs, albeit to variable degrees, i.e., RPS25 knock-down *in vitro* reduced poly-GA by 90%, poly-GP by 50%, and poly-GR by 30%^13^. Collectively, these and other studies^16^ suggest that a number of different factors play critical roles in RAN translation.

Although the present study uncovered a requirement for *eif-2D*/eIF2D in RAN translation, this protein has additional functions besides translation initiation. For example, eIF2D was implicated in recycling 40S ribosomal subunits^51, 52, 53^, which enabled ribosomes and mRNAs to participate in multiple rounds of translation, leading to efficient protein expression. It is thus possible that genetic removal of *eif-2D* in our worm models caused inefficient DPR production by blocking recycling 40S ribosomal subunits.

### RAN translation occurs in *C. elegans* and offers the opportunity to discover conserved regulators

Two previous studies reported DPR-associated toxicity in *C. elegans* by using either a codon-optimized construct that carries no G4C2 repeats, or a construct that carries 66 G4C2 repeats^22, 46^. The latter study strongly indicated that repeat associated non-AUG (RAN) translation occurs in the worm. However, the non-canonical (non-AUG) initiating codon and translation initiation factor remained elusive. By generating *C. elegans* animals that carry 75 G4C2 repeats flanked by intronic *C9orf72* sequences and subsequently blocking poly-GA expression by mutating a non-canonical initiation codon (CUG) upstream of the repeats, we advanced the molecular understanding of how RAN translation occurs in the worm. This finding in *C. elegans*, together with previous reports in yeast and flies ^16, 22^, strongly suggest that the molecular machinery required for RAN translation is evolutionary conserved, offering an opportunity to use simple model systems for the discovery of RAN regulators.

In summary, we leveraged the specific strengths of the *C. elegans* model and discovered that the non-canonical translation initiation factor *eif-2D/eIF2D* is required for RAN translation of expanded G4C2 repeats in the *C9orf72* gene. Our *in vitro* findings argue for a conserved role of *eif-2D/eIF2D* in RAN translation, potentially offering new directions for *C9orf72*-associated ALS and FTD therapy. In fact, because RAN translation of other nucleotide repeat expansions (not G4C2) occurs in several neurodegenerative diseases^4^, *eIF2D* may also prove to be a promising therapeutic target in these disorders. Determining how broadly eIF2D influences RAN translation is thus an important future endeavor.

## METHODS

### *C. elegans* strains

Worms were grown at 15°C, 20°C or 25°C on nematode growth media (NGM) plates supplied with *E. coli* OP50 as food source^54^. The mutant strains used in this study are *eif-2A (gk358198)* and *eif-2D (gk904876)*. The PCR primers used to genotype these mutant strains are shown in Supplementary Table 1. In addition, we used animals carrying the *vsIs48 [unc-17::gfp]* and *otIs437 [Punc-3::eGFP::rab-3cDNA::unc-10_3′UTR, + ttx-3::mCherry*] transgenes.

### Generation of C9 ^ubi^ and C9 ^neuro^ transgenic animals

The template for the generation of the plasmids used in the C9 ^*ubi*^, ΔC9 ^*ubi*,^ and UAG ^*ubi*^ animals have been described in an *in vitro* study^20^. In C9 ^*ubi*^ animals, the plasmid has 75 G4C2 repeats with 113 nucleotides 5’ and 99 nucleotides 3’ of intronic sequences that normally flank the G4C2 repeats in the human *C9orf72* gene; this sequence is followed by nanoluciferase (nLuc) reporter in the reading frame of poly-GA, which is followed by the *unc54* 3’ UTR. In ΔC9 ^*ubi*^ animals, the construct is identical but lacks the G4C2 repeats. In UAG ^ubi^ animals, the construct is identical but the non-canonical translation initiating codon CUG is mutated to UAG. In C9 ^*ubi*^, ΔC9 ^*ubi*^, UAG ^*ubi*^ animals, the following promoter sequence of the *snb-1* gene was used: cagttcgggtatctcagcaaccatgacgtgtgttcaagtatttcattttttctccctttctaggaaaattcaagttgaatttttattcattttgaaacttcctctcatctctcgaagatatgcctcgggcttcaaattcgtagtccacccgttgcacgtaacctgctgaaaacttccttgtattcagttctctcgtttcgttttgtctgcaaaagaatttagctccatttgtgtgaaatcgagcaaaatcatgcggctagagaaaataagaaaaaaaaacgaaagagaactggagcgtgtgcatgggacattctcctttcttcgccttattcttcgtctcccattattagtttcgcctcccacaattctggctgcaaagcaataaaaagtggcttcgttatttgtctggaatgcgtcccactgtatgtgttgtcttctatttaatacaagttataacctccactcgcttttttttttattttttgactgcctcttggtaaaaacgtaatcatacaacagcagtgaaaaccagttttttgaaaaccgtcccgtcattttttttctatttatcatttcaatttatttattaatcgat gattgaaagtgaatggatgacggtcatgaccgattatcgattatcccgaaatagagatgcgcgtaggtcataatgcccagtacgcaaaatgttttatcggtgtttgcacagatttcgcaacatctctcattgaatttccattcatcgcttcgtcatctgaccccatttcttattttttcatccttttccctgttctcatcgttccttactattttcctaatttcagaacatcgcgattttataatttcgttaaatattcgtaatcccgttatacaaaaatagctaaattttctagtcgttctcgtttttgagagggcactttagtccgtcatcgtgtcgcttgtcgtgctcaatttttcatgcataaatgggcgtcgccgtcccccctgtcgttttcttcctttacctcactttccagttctgaattccgatacgaatttttaaatttttctaactcgcttcatttcagggaacagccggataagaccatcttga cgac.

In C9 ^*neuro*^, ΔC9 ^*neuro*^, UAG ^*neuro*^ animals, the following promoter sequence of the *unc-11* gene was used: ctgtgtctctccgtcctatccactatctttcatccagattcccggttttcccgagaaaaagtgagagagagaagaggctactgcgcggcgacactggctgcggacaccctcactcctctacttgtctccatcgattttcgctggttcgtcgaagcctcgtcgccaacacaacttcttcttctttttcgtcctcttcttttctgccacacgcttgcacagcttcttcttctcattttacgctgtctacttttttcggaggggccatacccccacacacagctccttttattt gg.

The *unc54* 3’ UTR was cloned into the XhoI-SacI site of the plasmids using the primer sets shown in Supplementary Table 1. **Fig. 1A** shows the constructs with the *snb-1* promoter and the upstream CUG, which is the translation initiation codon for poly-GA (see Results).

The C9, UAG, or ΔC9 plasmids (100 ng/μl), along with a myo-2:GFP plasmid (3 ng/μl), and pBlueScript plasmid (20 ng/μl) were microinjected into the *C. elegans* gonad. Multiple transgenic lines per construct were established. The resulting C9 ^*ubi*^, ΔC9 ^ubi^ and UAG ^*ubi*^ animals, as well as the resulting C9 ^*neuro*^, ΔC9 ^*neuro*^ and UAG ^*neuro*^ animals, carried these transgenes as extrachromosomal arrays. The exact extrachromosomal lines used in each experiment are indicated in Figure legends. The following extrachromosomal lines were generated for this study: *kasEx153 [Psnb-1::C9 + myo-2::GFP line 3], kasEx154 [Psnb-1:: ΔC9 + myo-2::GFP line 1], kasEx155 [Psnb-1::UAG + myo-2::GFP line 1], kasEx156 [Punc-11::C9 + myo-2::GFP line 2.1], kasEx157 [Punc-11::C9; myo-2::GFP line 11.1], kasEx158 [Punc-11:: ΔC9 + myo-2::GFP line 3.1], kasEx159 [Punc-11:: ΔC9 + myo-2::GFP line 4.2], kasEx160 [Punc-11::UAG + myo-2::GFP line 4.1], and kasEx161 [Punc-11::UAG + myo-2::GFP line 8.1]*. For the C9 ^ubi^ and ΔC9 ^*ubi*^ extrachromosomal lines, we also established integrated versions into the *C. elegans* genome: *kasIs6 [Psnb-1::C9 + myo-2::GFP line 1.9], kasIs7 [Psnb-1::C9 + myo-2::GFP line 12.3], kasIs8 [Psnb-1:: ΔC9 + myo-2::GFP line 2.3]*, and *kasIs9 [Psnb-1:: ΔC9 + myo-2::GFP line 7.7]*.

### Lifespan assay

The lifespan assay was performed as previously described^22^. Worms were transferred at the L4 larval stage to NGM plates containing 50 μM 5-fluoro-2’deoxyuridine (FUDR, Sigma). The day of the timed egg laying was considered day 0 for lifespan analysis.

### Microscopy

Worms were anesthetized using 100mM of sodium azide (NaN_3_) and mounted on 5% agarose on glass slides. Images were taken using an automated fluorescence microscope (Zeiss, Axio Imager Z2). Acquisition of several z-stack images (each ~1 μm) Zeiss Axiocam 503 mono using the ZEN software (Version 2.3.69.1000, Blue edition). Representative images are shown following max-projection of 2-5 μm Z-stacks using the maximum intensity projection type. Image reconstruction was performed using Image J software^55^.

### Immunocytochemistry

C9 ^*ubi*^ and ΔC9 ^*ubi*^ animals were grown at 20 °C on nematode growth media (NGM). We followed an immunocytochemical staining procedure described previously^56^. In brief, worms were prepared for staining following the freeze-crack procedure and they were subsequently fixed in ice-cold acetone (5 min) and ice-cold methanol (5 min). Worms were transferred using a Pasteur pipette from slides to a 50mL conical tube that contained 40mL 1XPBS. Following a brief centrifugation (2min, 3000rpm), worms were pelleted and 1xPBS was removed. Next, the worms were incubated with 300 μl of blocking solution (1XPBS, 0.2% Gelatin, 0.25% Triton) for 30min at room temperature (rolling agitation). Following removal of the blocking solution, worms were incubated over-night with a mouse anti-poly-GA antibody [1:25 in PGT solution, EMD Millipore]. Next, the primary antibody solution was removed and worms were washed five times with washing solution (1XPBS, 0.25% Triton). Worms were incubated with a Alexa Fluor 594 goat anti-mouse IgG secondary antibody (1:1000 in PGT solution, A-11020, Molecular Probes) for 3 hours at room temperature. Following 5 washes, worms were mounted on a glass slide and examined at an automated fluorescence microscope (Zeiss, AXIO Imager Z1 Stand).

### Worm tracking

Worms were maintained as mixed stage populations by chunking on NGM plates with *E. coli* OP50 as the food source. Worms were bleached and the eggs were allowed to hatch in M9 buffer to arrest as L1 larvae. L1 larvae were refed on OP50 and allowed to grow to day 2 or adulthood. On the day of tracking, five worms were picked from the incubated plates to each of the imaging plates (see below) and allowed to habituate for 30 minutes before recording for 15 minutes. Imaging plates are 35 mm plates with 3.5 mL of low-peptone (0.013% Difco Bacto) NGM agar (2% Bio/Agar, BioGene) to limit bacteria growth. Plates are stored at 4°C for at least two days before use. Imaging plates are seeded with 50μl of a 1:10 dilution of OP50 in M9 the day before tracking and left to dry overnight with the lid on at room temperature.

### Behavioral feature extraction and analysis

All videos were analyzed using Tierpsy Tracker to extract each worm’s position and posture over time^35^. These postural data were then converted into a set of behavioral features selected from a large set of features as previously described^35^. For each strain comparison, we performed unpaired two-sample t-tests independently for each feature. The false discovery rate (FDR) was controlled at 5% across all strain and feature comparisons using the Benjamini Yekutieli procedure. The p-value threshold for this analysis (FDR 5%) is 0.0014.

### In vitro studies

Bicistronic plasmids were constructed with firefly luciferase (fLuc) in the first cistron and C9 in the second cistron, as previously published^20^.

shRNA plasmids against eIF2D were prepared using previously published methods^20^. In brief, oligonucleotides with an siRNA sequence (Supplementary Table 1) were cloned into the BamHI and Hind III sites of psilencer 2.1-U6 neo vector (Thermo Fisher Scientific) according to the manufacturer’s protocol. The control shRNA vector was provided by this vector kit.

For Western blotting (see below), HEK293 cells were planted in 6-well plates at 2 x 10^5^ per well and cotransfected with 0.5 μg bicistronic plasmids along with either 2.5 μg control shRNA or anti-eIF2D shRNA using Lipofectamine LTX (Thermo Fisher Scientific). For luciferase assays (see below), the cells were plated in 24-well plates at 5 x 10^4^ per well and cotransfected with 0.1 μg bicistronic plasmids along with either 0.5 μg control shRNA or anti-eIF2D shRNA using Lipofectamine LTX (Thermo Fisher Scientific). The cells were cultured for 48h in DMEM supplemented with 10% FBS, 2 mM L-Glutamine, 100 U/ml Penicillin and 100 μg/ml Streptomycin, and then processed.

### Western blotting

Lysates of worms were prepared by sonication in buffer consisting of 4% urea and 0.5% SDS with 1 x Halt^™^ Protease and Phosphatase inhibitor Cocktail (Thermo Fisher Scientific). HEK293 cell lysates were prepared using RIPA buffer consisting of 150 mM NaCl, 1% Nonidet P-40, 0.5% sodium deoxycholate, 0.1% SDS, 25 mM Tris-HCl (pH7.4). These lysates were subjected to electrophoresis on 4-20% SDS polyacrylamide gels (Mini-PROTEAN TGX Gels, BIO-RAD), and then transferred to Amersham Hybond P 0.45 □ μm PVDF membranes (GE Healthcare). The membrane was blocked with 1% non-fat skim milk in Tris-buffered saline containing 0.05% Tween-20 for 1□h at room temperature, and then incubated overnight at 4□°C with primary antibodies against poly-GA (1:1000, MABN889, EMD Millipore), fLuc (1:1000, G7451, Promega), eIF2D (1:2000, ab108218, Abcam) or tubulin (1:5000, YL1/2, Abcam). Following washing, the membrane was incubated for 1□h at room temperature with anti-mouse (1:5000, GE Healthcare), anti-rabbit (1:5000, GE Healthcare) or anti-rat horseradish peroxidase–conjugated secondary antibodies (1:1000, Cell Signaling Technology). The signal was detected using SuperSignal West Dura Extended Duration Substrate (Thermo Fisher Scientific) and analyzed using ChemiDoc MP Imaging System (BIO-RAD).

### Luciferase assay

The cells were lysed with 1x passive lysis buffer (Promega). Levels of nLuc and fLuc were assessed using Nano-Glo Dual-Luciferase Reporter assay system (Promega) and a Wallac 1420 VICTOR 3 V luminometer (Perkin Elmer) according to the manufacturer’s protocol.

### RT-PCR

Total RNA was extracted using TRIzol (Thermo Fisher Scientific) according to the manufacturer’s protocol, treated with TURBO DNase (Thermo Fisher Scientific) for 30□min at 37ū°C, and RNA was then isolated using RNeasy Mini kit (Qiagen). cDNA was generated using SuperScript IV First-Strand Synthesis System (Thermo Fisher Scientific). Quantitative RT-PCR was performed using Power SYBR Green PCR Master Mix (Thermo Fisher Scientific) and primer sets (Supplementary Table 1) in a CFX96 Real-Time System (Bio-Rad).

### ELISA

The worm pellets were suspended in cold Co-IP buffer consisting of 50 mM Tris-HCl, pH 7.4, 300 mM NaCl, 5mM EDTA, 1% Triton-X 100, 2% sodium dodecyl sulfate, 1 x Halt^™^ Protease and Phosphatase Inhibitor Cocktail and sonicated with 10 cycles of 0.5 second pulse/0.5 second pause. Protein lysates were cleared by centrifugation at 16,000 g for 20 min at 4°C. The protein concentration was determined using a BCA protein assay kit (Thermo Fisher Scientific). Lysates were diluted to the same concentration using Co-IP buffer, and the levels of poly-GP in lysates were measured with a Meso Scale Discovery-based immunoassay as described previously^57^.

### Statistical analysis

Statistical analysis was performed by an unpaired t-test, one-way ANOVA with Dunnett’s multiple comparisons test, and two-way ANOVA with the Šídák multiple comparison test using GraphPad Prism version 8.2.1. P values for the worm lifespan assay were calculated by the Mantel-Cox log-rank test using GraphPad Prism 8.2.1. A P-value of <0.05 was considered significant. The data are presented as mean ± standard error of the mean (SEM).

## Acknowledgements

We thank the *Caenorhabditis* Genetics Center (CGC), which is funded by NIH Office of Research Infrastructure Programs (P40 OD010440), for providing strains. This work was supported by grants from the Lohengrin Foundation and the Association for Frontotemporal Degeneration (AFTD).

## Author Contributions

Study design: Y.S., P.K., R.P.R. Literature research: Y.S., P.K., R.P.R. Experimental studies: Y.S., J.A., P.K., T.F.G., P.I. Data analysis/interpretation: Y.S., A.E.X.B., P.K., R.P.R. Statistical analysis: Y.S., A.E.X.B. Manuscript preparation: Y.S., P.K., R.P.R. Manuscript editing: Y.S., P.K., R.P.R.

## SUPPLEMENTARY FIGURE LEGENDS

**Supplementary Figure 1. Assays to detect poly-GR in worm lysates.** (A) The lysates from ΔC9^*ubi*^ and C9^*ubi*^ worms were processed for Western blotting and immunostained with poly-GR and a-tubulin antibodies. The lysate from HEK293 cells transfected with GR^AUG^(G4C2_75_-nLuc) plasmid, in which AUG start codon was inserted before G4C2 repeats in the frame of poly-GR, was used as a positive control. (B) The lysates from ΔC9*^neuro^*, C9^*neuro*^, and UAG^*neuro*^ worms were processed for GR ELISA (n = 4, mean ± s.e.m.).

**Supplementary Figure 2. Genetic removal of *eif-2D/eIF2D* does not affect expression of an AUG-initiated green fluorescent protein (GFP) transgene.** The ΔC9 ^*ubi*^ transgene also carries a pharyngeal *gfp* marker (*myo-2::gfp*), but its expression is not affected in *eif-2D (gk904876)* animals. Representative images of larval stage 4 (L4) worms are shown. Differential interference contrast (DIC) images of the worm head are shown on top. Dashed white line demarcates the pharynx. Fluorescent images are shown below in the GFP channel. N = 10.

**Supplementary Figure 3. Defects in velocity and body curvature are ameliorated in *C9 ^ubi^* animals upon genetic removal of *eif-2D/eIF2D*.**

Examples of seven additional locomotion features significantly affected in *C9 ^ubi^* animals. Genetic removal of *eif-2D* ameliorated the locomotion defects of *C9 ^ubi^* animals [*eif-2D (-); C9 ^ubi^*]. Tracking analysis was performed at Day 2 adult animals (N = 12). The p-value threshold for this analysis (FDR 5%) is 0.0014 (see Methods). Panel A shows features related to animal velocity. *d ang vel head base w forward abs 90^th^*: 90th percentile of the derivative of the absolute value of the angular velocity of the base of the head, while the worm is moving forwards.

*rel to hips radial vel tail tip w forward IQR*: interquartile range of the radial velocity of the tip of the head relative to the centroid of the hips points, while the worm is moving forwards.

*d rel to tail base radial vel tail tip 50^th^*: 50th percentile of the derivative of the radial velocity of the tip of the tail relative to the centroid of the tail base points.

*d rel to body radial vel tail tip w backward 10^th^*: 10th percentile of the derivative of radial velocity of the tip of the tail relative to the centroid of the midbody points, while the worm is moving backwards. Panel B shows a feature related to body curvature. *d curvature mean hips abs IQR*: interquartile range of the derivative of the absolute value of the mean curvature of the hips.

## SUPPLEMENTARY INFORMATION

**Supplementary Table 1:** Sequences of oligonucleotides used in this paper.

