## Supplementary Figures for "A *C. elegans* model of *C9orf72*-associated ALS/FTD uncovers a conserved role for eIF2D in RAN translation"

Supplementary Figure 1

A

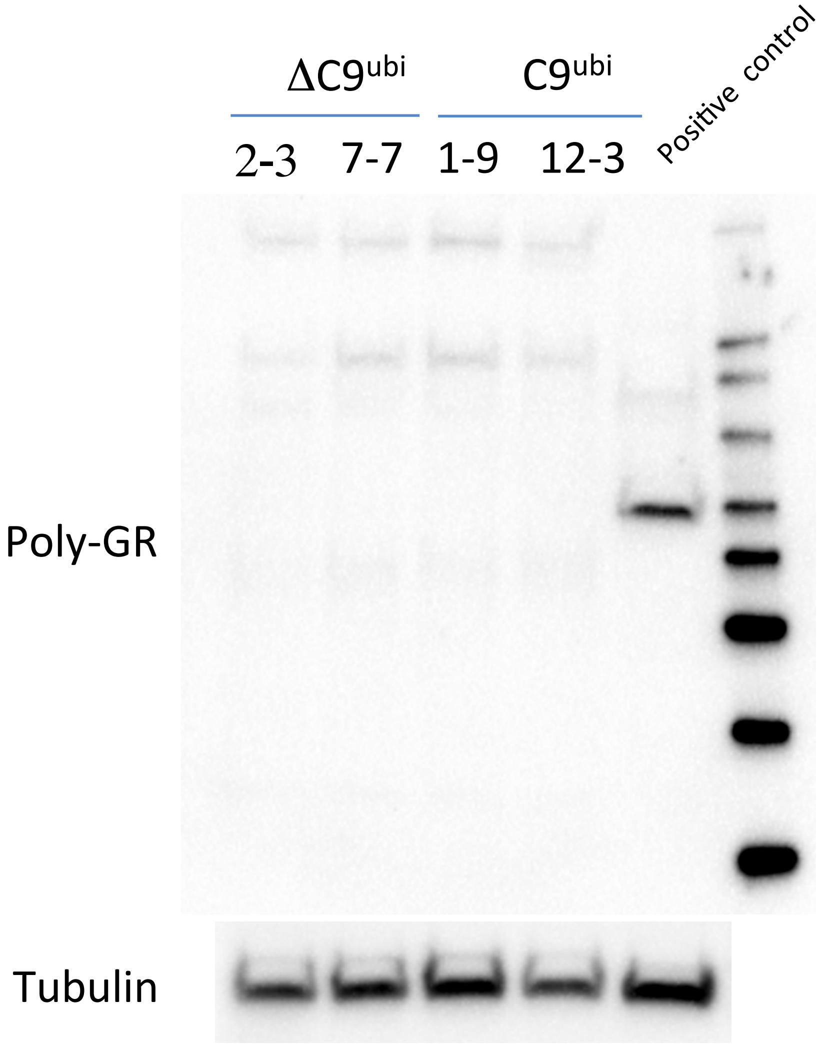

B

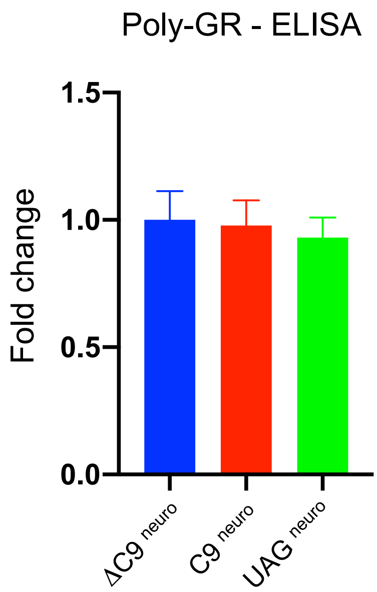

### Supplementary Figure 2

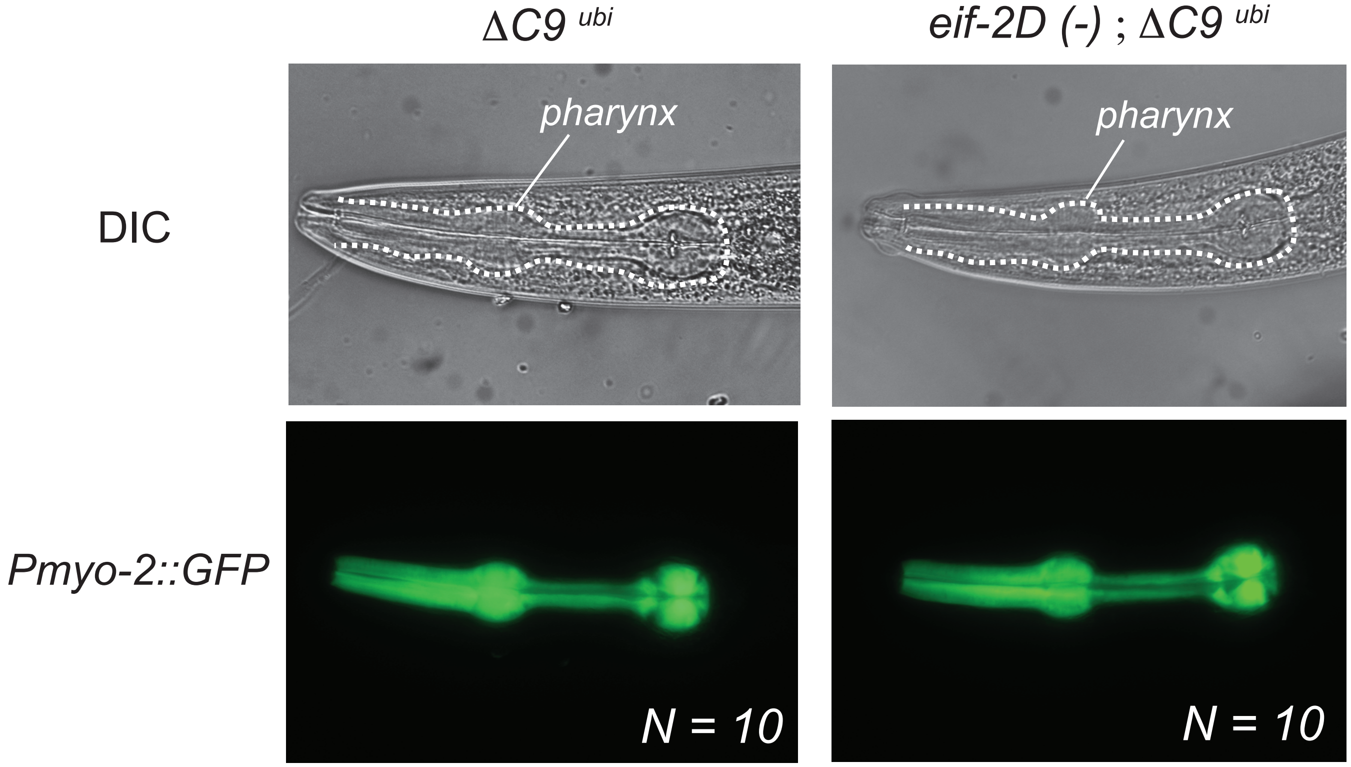

### Supplementary Figure 3

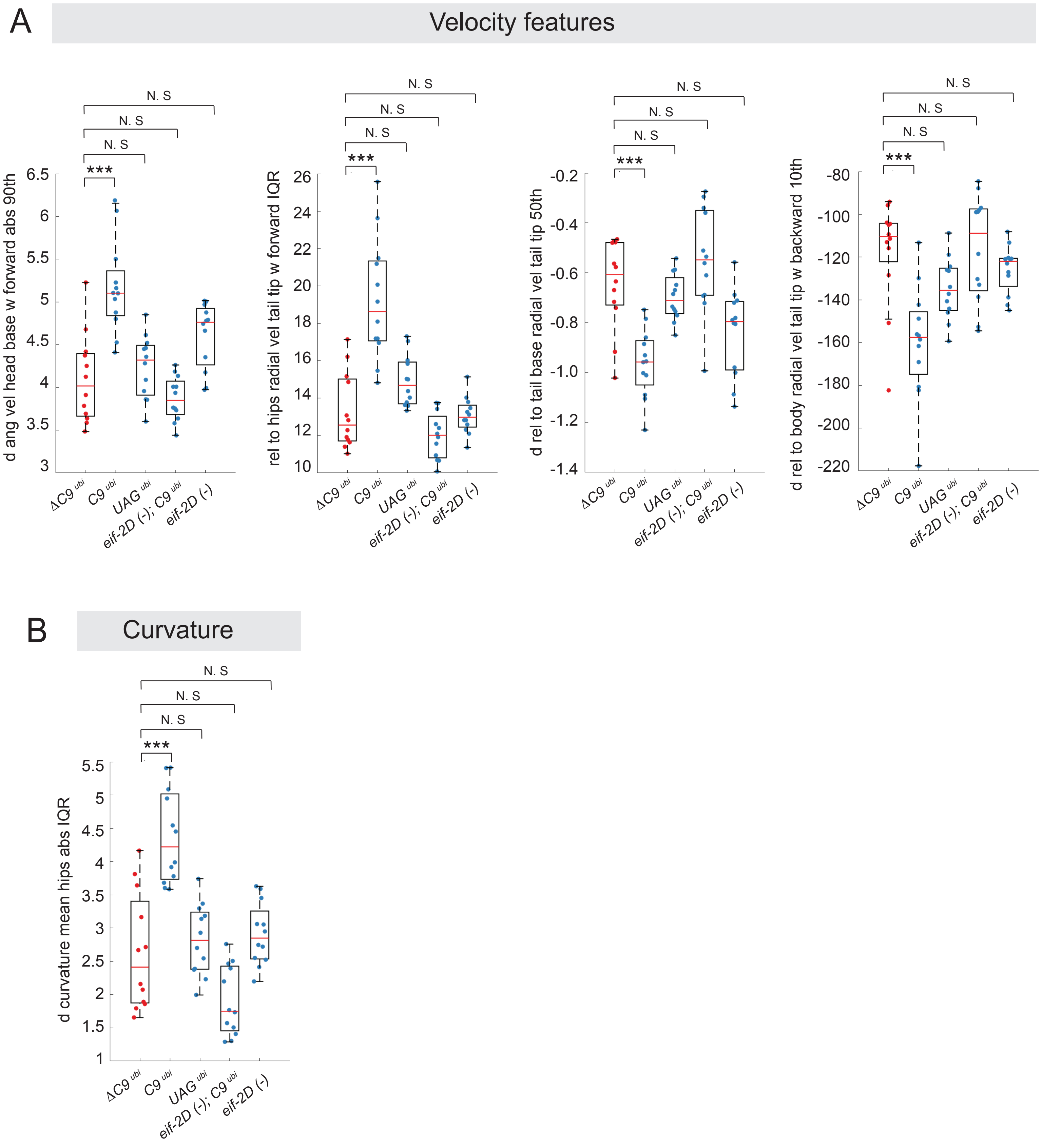

Supplementary Table 1 Sequences of oligonucleotides

| Description | Sequence (5'->3') |
|---|---|
| Ce-eIF2A-F | aaggaaaaaagctagctgggcagtttacggaccaaa |
| Ce-eIF2A-R | aaggaaaaaacttaaggaagttggtcgcacccttct |
| Ce-eIF2D-F | aaggaaaaaagctagcaccgtttacagtcaagaaaaataca |
| Ce-eIF2D-R | aaggaaaaaacttaaggtgtagacattcatctgggttcct |
| Hs-eIF2D-shRNA-F | gatccgggacgacaactggacataaagttcaagagactttatgtccagttgtcgtcctttttggaaa |
| Hs-eIF2D-shRNA-R | agcttttccaaaaaaggacgacaactggacataaagtctcttgaactttatgtccagttgtcgtcccg |
| Ce-qPCR-Act-1-F | acgacgagtccggcccatcc |
| Ce-qPCR-Act-1-R | gaaagctggtggtgacgatggtt |
| C9-qPCR-C9-F | gaaacaaccgcagcctgtag |
| C9-qPCR-C9-R | tagcgcgcgactcctga |
| Ce-pSnb-1-F | aaacccactagtcagttcgggtatctcagcaa |
| Ce-pSnb-1-R | acagggaagcttgtcgtcaagatggtcttatc |
| Ce-pUnc11c-F | aaacccactagtctgtgtctctccgtcctatc |
| Ce-pUnc11c-R | aaacccaagcttccaaataaaaggagctgtgt |
| Ce-Unc54-F | aaacccctcgaggcgccggtcgctaccattac |
| Ce-Unc54-R | acaggggagctcggaaacagttatgtttggta |
